# Firing rate homeostasis occurs in the absence of neuronal activity-regulated transcription

**DOI:** 10.1101/675728

**Authors:** Kelsey M. Tyssowski, Katherine C. Letai, Samuel D. Rendall, Anastasia Nizhnik, Jesse M. Gray

**Affiliations:** Department of Genetics; Harvard Medical School; Boston, MA 02115

## Abstract

Despite dynamic inputs, neuronal circuits maintain relatively stable firing rates over long periods. This maintenance of firing rate, or firing rate homeostasis, is likely mediated by homeostatic mechanisms such as synaptic scaling and regulation of intrinsic excitability. Because some of these homeostatic mechanisms depend on transcription of activity-regulated genes, including *Arc* and *Homer1a*, we hypothesized that activity-regulated transcription would be required for firing rate homeostasis. Surprisingly, however, we found that cultured mouse cortical neurons grown on multi-electrode arrays homeostatically adapt their firing rates to persistent pharmacological stimulation even when activity-regulated transcription is disrupted. Specifically, we observed firing rate homeostasis *Arc* knock-out neurons, as well as knock-out neurons lacking activity-regulated transcription factors, AP1 and SRF. Firing rate homeostasis also occurred normally during acute pharmacological blockade of transcription. Thus, firing rate homeostasis in response to increased neuronal activity can occur in the absence of neuronal-activity-regulated transcription.

**SIGNIFICANCE STATEMENT:** Neuronal circuits maintain relatively stable firing rates even in the face of dynamic circuit inputs. Understanding the molecular mechanisms that enable this firing rate homeostasis could potentially provide insight into neuronal diseases that present with an imbalance of excitation and inhibition. However, the molecular mechanisms underlying firing rate homeostasis are largely unknown.

It has long been hypothesized that firing rate homeostasis relies upon neuronal activity-regulated transcription. For example, a 2012 review (PMID 22685679) proposed it, and a 2014 modeling approach established that transcription could theoretically both measure and control firing rate (PMID 24853940). Surprisingly, despite this prediction, we found that cortical neurons undergo firing rate homeostasis in the absence of activity-regulated transcription, indicating that firing rate homeostasis is controlled by non-transcriptional mechanisms.

## INTRODUCTION

Neuronal circuits maintain relatively stable firing rates even in the face of changes in circuit inputs that result from sensory stimuli and experience-dependent plasticity. For example, upon eliminating visual input to the mouse visual cortex via monocular deprivation, the firing rates of visual cortex neurons initially decrease, but over a period of 3-4 days, they return to the pre-deprived levels (Hengen et al., 2016, 2013), thus undergoing homeostasis. Firing rate homeostasis has also been observed in cultured neurons in response to chronic stimulation or chronic blockade of neuronal activity (Bateup et al., 2013; Burrone et al., 2003; Pozzi et al., 2013; Slomowitz et al., 2015; Turrigiano et al., 1998). Understanding the molecular mechanisms that enable firing rate homeostasis may provide insight into human diseases that present with an imbalance of excitation and inhibition in the brain, such as epilepsy or autism (Turrigiano, 2011). However, the molecular mechanisms underlying firing rate homeostasis are largely unknown.

We hypothesized that activity-regulated transcription would be required for firing rate homeostasis because it is required for several forms of homeostatic plasticity that could regulate firing rate. First, synaptic scaling, a multiplicative homeostatic change in the number of AMPA receptors at synapses (Turrigiano et al., 1998), is impaired by both acute blockade of transcription and by loss of specific activity-regulated genes (ARGs), including *Arc, Nptx1, Plk2*, and *Homer1a* (Chowdhury et al., 2006; Diering et al., 2017; Hu et al., 2010; Ibata et al., 2008; Schaukowitch et al., 2017; Seeburg et al., 2008; Shepherd et al., 2006). In addition, activity-regulated transcription also homeostatically regulates excitatory/inhibitory (E/I) balance, or the relative strengths and numbers of excitatory and inhibitory synapses. Acute blockade of transcription following stimulation impairs homeostatic decreases in excitatory synapse number (Goold and Nicoll, 2010), and ARGs, such as *Nptx2, Igf1, Bdnf*, and *Npas4*, homeostatically regulate inhibitory synapse number (Bloodgood et al., 2013; Chang et al., 2010; Gray and Spiegel, 2019; Hartzell et al., 2018; Spiegel et al., 2014). Finally, transcription mediates homeostatic changes in intrinsic excitability, that is, the composition of ion channels in the neuronal membrane that determine a neuron’s likelihood of firing an action potential upon stimulation (Turrigiano, 2011). In the *Drosophila* neuromuscular junction, the transcription factor Kruppel is required for homeostatic alterations in intrinsic excitability and in firing rate, likely due to its regulation of potassium channels (Kulik et al., 2019; Parrish et al., 2014). While it is unclear whether Kruppel regulates activity-dependent or basal transcription, the mammalian ARG program has an enrichment for potassium channels (Cho et al., 2016), suggesting that activity-dependent potassium channel transcription may homeostatically change intrinsic excitability. Importantly, each of these transcription-dependent forms of homeostatic plasticity—synaptic scaling, changing E/I balance, and changing intrinsic excitability—can alter firing rates and are thus leading candidate mechanisms underlying firing rate homeostasis (Turrigiano, 2012). Consistent with this idea, synaptic scaling and homeostatic changes in intrinsic excitability both occur in neurons undergoing firing rate homeostasis (Bateup et al., 2013; Burrone et al., 2003; Hengen et al., 2013; Slomowitz et al., 2015; Turrigiano et al., 1998). Thus, we reasoned that transcription might regulate firing rate homeostasis through one or more of these forms of homeostatic plasticity thought to underlie firing rate homeostasis.

Another reason that we considered ARG transcription to be a promising candidate regulator of firing rate homeostasis is that ARG transcription and firing rate homeostasis both occur over a period of several hours following chronic stimulation or activity blockade (Bateup et al., 2013; Hengen et al., 2013; Schaukowitch et al., 2017; Slomowitz et al., 2015; Tyssowski et al., 2018; Yap and Greenberg, 2018). Furthermore, a model that takes into account the kinetics of ARG induction and the mechanisms mediating their induction suggests that activity-regulated transcription of ion channels could regulate firing rate homeostasis (O’Leary et al., 2014). Therefore, activity-regulated transcription is a strong candidate regulator of firing rate homeostasis based on both the composition of the ARG program and the kinetics of ARG induction.

Here we test the role of activity-regulated transcription in firing rate homeostasis that occurs in response to increases in neuronal activity. We observe firing rate homeostasis in individual cultured cortical neurons plated on multi-electrode arrays (MEAs). Following increases in firing rate induced by GABA_A_ blockade with picrotoxin (PTX), neurons return to their pre-stimulation firing rates over a period of 30 hours. Surprisingly, we find using knock-out mice that neurons lacking either the activity-regulated gene *Arc* or the activity-regulated transcription factors SRF or AP1 can still undergo firing rate homeostasis. Furthermore, neurons are also able to achieve firing rate homeostasis in the presence of an acute pharmacological blockade of activity-regulated transcription. Thus, firing rate homeostasis still occurs in the absence of activity-regulated transcription.

## MATERIALS AND METHODS

### Animal Care

All animal care and experimental procedures were approved by the Institutional Animal Care and Use Committees at Harvard Medical School. Animals were housed with standard mouse chow and water provided *ad libitum*.

### Mouse Strains

Wildtype CD-1 litters were acquired from Charles River (Woburn, MA). The Arc KO strain (007662, The Jackson Laboratory, Farmington, CT) expresses a destabilized form of GFP in the place of Arc under the Arc promoter (Wang et al., 2006). The SRF KO strain (Jackson 006658) contains loxP sites flanking the promoter and exon 1 sequences of SRF. The AP1 KO strain is a triple-transgenic acquired from the Greenberg Lab (Vierbuchen et al., 2017), contains loxP sites at three AP1 subunits: Fos (Fleischmann et al., 2003), Fosb (created in Vierbuchen et al., 2017), and Junb (Kenner et al., 2004).

### Neuronal Culture

Cortical neurons were dissected from pups of mixed sex on postnatal days 0-2 and dissociated with papain ((L)(S)003126, Worthington, Lakewood, NJ). Neurons were plated on standard plastic plates coated with poly-ornithine (30mg/mL, Sigma Aldrich) in water, or on Lumos MEA plates (Axion Biosystems, Atlanta, GA) or Lumos OptiClear plates (Axion Biosystems) coated with poly-ornithine (30mg/mL) and 5ug/mL laminin (Gibco, Cambridge, MA). Neurons were cultured in BrainPhys media (Stem Cell Technologies), supplemented with SM1 (Stem Cell Technologies, Cambridge, MA), penicillin-streptomycin (Gibco), and fungizone (Gemini Bio, West Sacramento, CA). Cultures were incubated at 37°C in 5% CO_2_. Experiments were performed on neurons between DIV 15-28. For conditional knock-out experiments (SRF and AP1), neurons were treated with homemade AAV-CaMKII-GFP or AAV-CaMKII-Cre on DIV 3-4 for 3 days, and experiments were performed at least 14 days following viral treatment.

### Viral Production

AAV-CamKII-Cre and AAV CamKII-eGFP virus were made by transfecting HEK293T cells with the particles pHelper, RC1, RC2, and CamKII-Cre or CamKII-eGFP. Viral supernatant was collected on day 3 of transfection, by collecting viral supernatant after 3 freeze/thaw cycles at −80°C and 37°C. Cellular debris was removed by centrifugation at for 3220g for 15 minutes, and filtering through a 0.45um filter. Virus was stored at 4°C for up to 1 month.

### MEA Recordings

Recordings were made using the Maestro and MiddleMan from Axion Biosystems (version 1.0.0.0), along with Axion’s AxIS software (version 2.4.5). Lumos MEA plates have 48 wells, each containing 16 PEDOT electrodes in a 4×4 grid. Electrodes are 50 μm in diameter and spaced 350 μm apart. Neurons were kept at 37°C with 5% CO_2_ during recordings using the Axion Maestro system.

### Spike Sorting

Raw data was filtered in AxIS online using a 200 Hz Butterworth high-pass filter and a 3000 Hz Butterworth low-pass filter. Spikes were detected in AxIS online using peak detection with an adaptive threshold of 5.5 standard deviations from noise levels. To avoid detection of overlapping spikes, detection was prevented for 2.16 ms after each peak. Spikes were semi-automatically sorted offline in MATLAB using custom-written code clustering waveforms (vectors of voltage over time, detected by AxIS) in principal component space and clustered by fitting to Gaussian mixture models, with the number of clusters determined manually.

### Homeostasis Assay

In each experiment, the MEA plate was moved to the Maestro at least 2 hours before recording baseline firing rate, in order to exclude from the recording any activity changes resulting from moving the plate from the incubator to the machine. The baseline recording lasted 6 hours, and then PTX (2.5uM, in DMSO and water; Tocris, Minneapolis, MN) or DMSO (1:1000 in water) was added to individual wells and activity was recorded for an additional 30 hours. DMSO-treated neurons received twice the amount of DMSO as PTX-treated neurons. In ActD experiments, ActD (1ug/mL) was added to individual wells for 30 minutes before PTX or DMSO. All drugs remained in the cultures for the duration of the 30-hour experiment.

One replicate each for Arc KO, SRF cKO and AP1 cKO were treated with 1uM PTX instead of 2.5uM PTX. However, the post-stimulus fold induction of firing rate in this replicate was comparable to replicates receiving 2.5uM PTX treatment, so we included the replicate in our analysis.

While refining this assay, we tried stimulating activity with other drugs in place of PTX, with a range of success. We chose to focus on PTX because it most consistently caused an increase in firing rate and subsequent homeostasis of firing rate. The other drugs attempted, along with a brief description of their effects on activity, can be found in Table S1.

### MEA Data Analysis

Spike data was binned into 5 minute bins in MATLAB (Mathworks, Natick, MA) and then analyzed further in Python and R. Median firing rate was determined for each experimental condition by taking the median firing rate of all neurons (across multiple wells) at each time bin. Baseline firing rate, post-stimulation firing rate, and final firing rate were determined for each condition by calculating the average median firing rate in the last 3 hours of baseline, 30 minutes to 3 h 30 min after adding PTX, and the last 3 hours of the 30-hour PTX exposure (respectively).

We included in the analysis only units whose firing rates were stably above 0.001 Hz for at least 28 hours of the experiment, and whose post-stimulation firing rate was greater than baseline firing rate. PTX-treated units were excluded if their post-stimulation firing rate was not greater than baseline firing rate. Furthermore, if the post-stimulation firing rate of a single PTX-treated well was not at least 1.3 fold of baseline firing rate, that well’s response was considered too weak and it was excluded from analysis. An entire experiment was excluded if the median firing rate of any 5 hour period in the DMSO control condition was below 0.75 fold or above 1.3 fold of baseline firing rate, with the exception of a single AP1 cKO biological replicate for which we did not have a DMSO control.

To determine the length of time it took for median firing rates to achieve homeostasis, we first calculated a rolling standard deviation of the median firing rate over 6h bins. We then determined homeostasis firing rate, that is the firing rate at the time point after PTX addition where the rolling standard deviation was lowest. We said that the median had achieved homeostasis at the time point post-PTX when the median firing rate first reaches at least 102.5% of the homeostasis firing rate.

### Code Accessibility

Custom code for both spike sorting and analysis is available on GitHub at https://github.com/kletai/mea_analysis_homeostasis. The MATLAB portions of this analysis were conducted on the O2 High Performance Compute Cluster, supported by the Research Computing Group, at Harvard Medical School. See http://rc.hms.harvard.edu for more information.

### RNA Extraction and qRT-PCR

Samples were collected in Trizol (Ambion, Cambridge, MA) and stored at −80°C. The RNeasy Mini Kit (Qiagen, Germantown, MD) was used to extract RNA according to the manufacturer’s instructions, including in-column DNase treatment (Qiagen) cDNA was reverse transcribed using the High Capacity cDNA Reverse Transcription kit (Applied Biosystems, Foster City, CA). For qPCR, SsoFast Evagreen Supermix (BioRad, Hercules, CA) was used with the primers in Table S2.

### Immunocytochemistry

Neurons were grown on glass coverslips coated with 5ug/mL laminin and 30mg/mL polyornithine as above. At least 14 days following viral treatment, they were fixed with 4% PFA for 15 minutes. Neurons were then washed twice in PBS and blocked and permeablized for 30 min using 1%BSA in PBS + 0.25% Triton X-100 (BSA-PBST). Neurons were then incubated overnight at 4C in BSA-PBST and primary antibody. They were then washed 3 times with PBS and incubated for 1h at room temperature in secondary antibody (1:1000; ThermoFisher, Cambridge, MA; R37117). They were then washed once with PBS, incubated for 10 min with DAP1 (Roche, Indianapolis, IN; 10236276001) in PBS, and washed again with PBS. They were mounted onto coverslips with Fluoromount-G. Primary antibodies used were: FOSB (Abcam, Cambridge, MA; #11959, 1:250), JUNB (Santa Cruz, Dallas, TX; #8051X, 1:250).

### Western Blotting

Samples were collected in cold lysis buffer (1% Triton X-100, 50 mM HEPES, pH 7.4, 150 mM NaCl, 1.5 mM MgCl2, 1 mM EGTA, 10% glycerol, and freshly added phosphatase inhibitors from Roche Applied Science Cat. # 04906837001). Lysed neurons were mixed 3:1 with 4X sample buffer (40% glycerol, 8% SDS, 0.25M Tris-HCL, pH 6.8, 10% 2-mercaptoethanol) and boiled for 5 min. Samples were centrifuged at full speed for 3 min before loading on NuPage 4%–12% Bis-Tris Gels (Invitrogen, Carlsbad, CA). Gels were run at 140V for 55 min. We transferred onto nitrocellulose mem-branes using the BioRad transfer system at 114V for 1 hr and 7 min. Membranes were blocked in 5% milk-Tris-buffered saline + Triton X-100 (TBST) for 1 hr. They were treated with primary antibody in 5% milk-TBST for overnight at 4°C. To visualize protein, blots were incubated with secondary antibody in TBST in the dark for 45 min. Blots were imaged using a Li-Cor Odyssey. Primary antibodies used include: anti-SRF (Cell Signaling Technology, Danvers, MA; #5147, 1:1000), anti-ARC (Synaptic Systems, Goettingen, Germany; # 156003, 1:1000). Secondary antibodies used were: IDR dye 680 goat anti-mouse (Li-Cor, Lincoln, NE; 1:10000), IDR dye 800 goat anti-rabbit (Li-Cor, 1:10000).

### Statistical Analysis

Statistical analysis was performed in R with the tests listed in the text.

Binned data from the units used to make plots and perform statistical analysis is available in Tables S3-S8.

## RESULTS

### Neurons undergo firing rate homeostasis in response to PTX treatment

To assess firing rate homeostasis, we measured the firing rates of individual neurons over a period of ~30 hours in the presence of a pharmacological perturbation that increased neuronal activity. We cultured dissociated neonatal mouse cortical neurons on microelectrode arrays (MEAs) and stimulated them with the GABA_A_ receptor antagonist PTX (2.5uM). In addition to PTX, we also tested other pharmacological stimuli but found that PTX was the most reliable for inducing increases in firing rate that were followed by firing rate homeostasis (see Methods, Table S1). To obtain the firing rates of individual units that likely represent single neurons, we spike-sorted the action potential wave forms obtained from MEA recordings (see Methods). In each experiment, we first recorded baseline neuronal firing rates over a period of 3 hours. We then stimulated with PTX, which induces an initial 2-fold median change in firing rate (Figure 1A, S1A). Over the next ~20 hours, the median neuronal firing rate returned to baseline values (Figure 1A, S1B). This adaptation of median firing rate could occur even if the firing rates of many neurons in the culture do not adapt. We therefore asked whether the distribution of firing rates was altered by PTX treatment. While PTX treatment initially shifts the distribution of firing rates (Figure 1B), the firing rate distribution post-homeostasis (i.e., the last three hours of recording) is not significantly different from the distribution during the baseline measurement (Figure 1C), suggesting that the distribution of firing rates in the circuit also homeostatically adapts. Therefore, we observe firing rate homeostasis in response to PTX stimulation.

**Figure 1.**
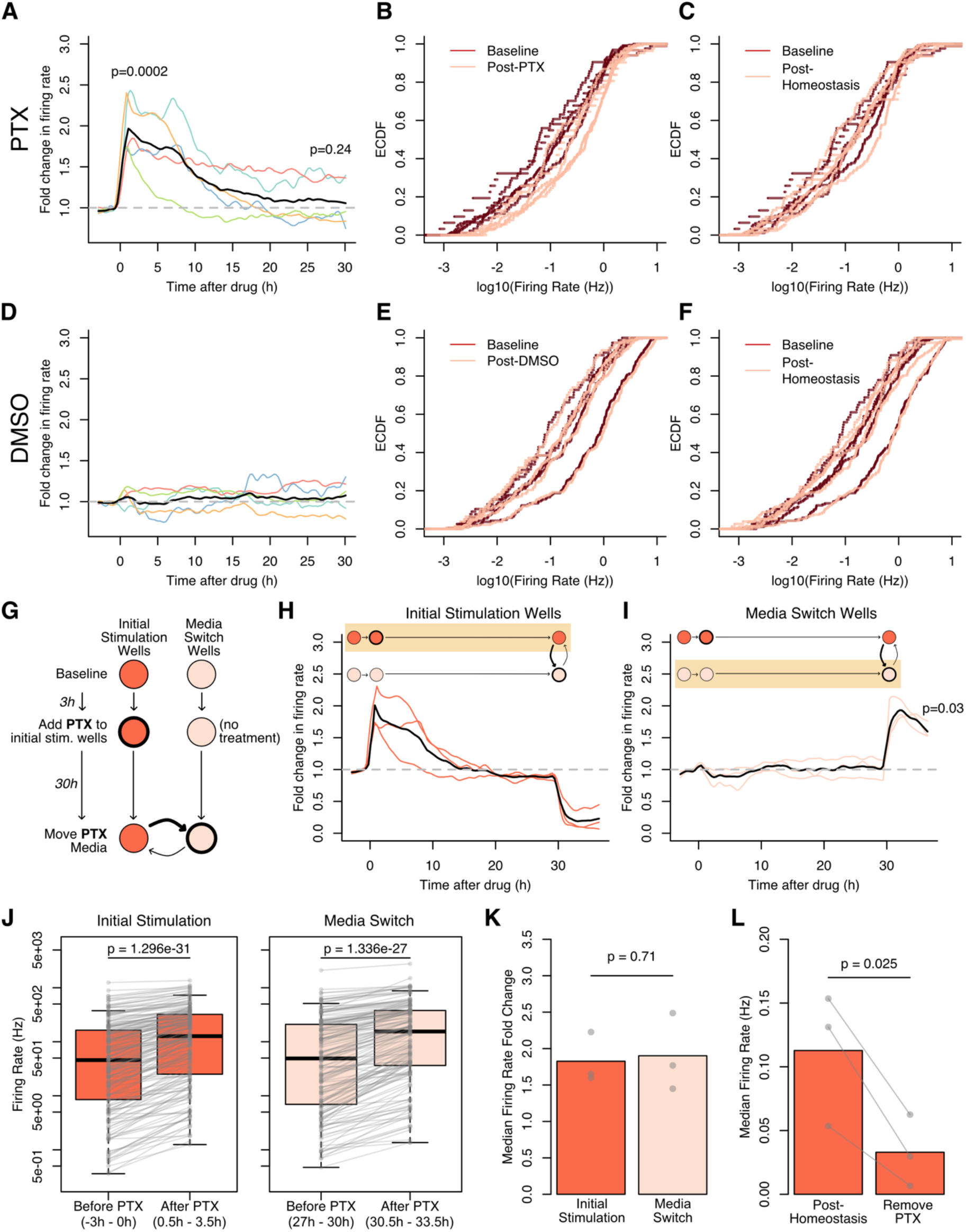
Cultured neurons homeostatically adapt their firing rates in response to GABA_A_ blockade. (A) Firing rate homeostasis following PTX stimulation. PTX (2.5uM) was added at time 0. Colored lines are medians of multiple neurons from each of n=5 biological replicates (31-182 units/replicate). The black line represents the mean of these medians. There is no significant fold change in firing rate from the first to last three hours (−3h to 0h, vs. 27-30h), p = 0.47, paired t-test on log(fold change in firing rate). (B) Cumulative distributions of mean firing rates of each unit before and after adding PTX (−3h to 0h, baseline; 0.5-3.5h, post-PTX). For each biological replicate (same data as in A), one baseline and one post-PTX curve is shown. For all replicates except one, the distribution of firing rates is significantly different between the baseline and post-PTX (Kolmogorov-Smirnov test, p<0.05). (C) Cumulative distributions of mean firing rates of each unit before adding PTX (baseline, as in B) and in the last three hours of the recording (post-homeostasis). For all biological replicates (same data as in A), the distribution of firing rates is not significantly different between the baseline and post-PTX (Kolmogorov-Smirnov test, p>0.05). (D) Same as (A) but for neurons treated with DMSO rather than PTX. n=5 biological replicates (77-187 units/replicate). (E) Same as (B) but for neurons treated with DMSO rather than PTX. For all replicates (same data as in B), the distribution of firing rates is not significantly different between the baseline and post-PTX (Kolmogorov-Smirnov test, p>0.05). (F) Same as (C) but for neurons treated with DMSO rather than PTX. For all replicates (same data as in B), the distribution of firing rates is not significantly different between the baseline and post-PTX (Kolmogorov-Smirnov test, p>0.05). (G) Schematic of the media switch experiment to test PTX potency. Half of the wells were stimulated with PTX at time 0h. At time 30h, the media from the stimulated wells (initial stimulation) was switched with those from unstimulated wells (media switch). (H) The fold change in firing rate over the course of the media switch assay in the wells initially stimulated with PTX, n=3 biological replicates (84-182 units/replicate). PTX (2.5uM) was added at 0h and media swapped (i.e., PTX removed) at 30h. Colored lines are medians from individual replicates. The black line represents the mean of these medians. (I) The fold change in firing rate over the course of the media switch assay in the wells that received PTX upon media switch, n=3 biological replicates (71-159 units/replicate). PTX-containing media from initially-stimulated wells was swapped in at 30h. Colored lines are medians from individual replicates, and the black line is the mean of these medians. P-value from a paired t-test on log(fold change) 3h before media switch (27h-30h) vs 3h after media switch (30-33h). (J) A representative replicate showing the change in firing rate upon addition of PTX in wells initially stimulated with PTX (left, n=182 units) and wells that received PTX upon media switch (right, n=159 units). Each point represents the mean firing rate of an individual neuron over a three hour period. P-values from a paired rank-sum test. p<10^−12^ for all replicates. (K) Comparison of the median fold changes in firing rate upon initial PTX addition and media switch. Bars represent the mean of the median fold changes. Dots represent the median fold change in each replicate. P-value from a paired t-test on the log fold changes. (L) Median firing rates upon removal of PTX from the initially-stimulated wells. Bars represent the mean of the median fold change. Dots represent the median fold change in each replicate. P-value from a paired, one-sided t-test on log(median firing rates).

We next asked whether the apparent firing rate homeostasis that we observed is genuine or due to potential artifacts of the assay. First, we asked whether the increase in neuronal activity that we observed following PTX treatment was due to GABA_A_ blockade or if it was a side effect of adding PTX (*e.g*., media perturbation). We found that neurons treated with a DMSO vehicle do not exhibit an increase in median firing rate or a change in the distribution of firing rates over the 30 hours following addition of the vehicle (Figure 1D-F, S1A,C), suggesting that the increase in neuronal activity upon stimulation is due to the PTX-mediated GABA_A_ block. Next, we asked whether the firing rate homeostasis that we observed can be explained by a decrease in PTX potency over course of the assay. We transferred PTX-containing media from PTX-treated neurons to untreated neurons at the end of a 30-hour experiment (Figure 1G). The addition of PTX-containing media increased the firing rates of the previously-untreated neurons to the same extent as the initial PTX treatment (Figure 1H-K, S1F-G), indicating that the PTX maintains its full potency throughout the experiment. In contrast, neurons newly treated with DMSO-containing media exhibited no such increase in firing rate (Figure S1E-F). Finally, we asked whether the firing rate homeostasis we observe is due to a desensitization of the GABA_A_ receptor to PTX. If the PTX-mediated GABA_A_ blockade were to persist throughout the end of the experiment, removal of PTX should induce an immediate increase in inhibition and decrease in firing rates. Indeed, we observed a decrease in firing rates upon removal of PTX post-homeostasis (Figure 1H,L, S1H), suggesting that the firing rate homeostasis we observe results from neurons lowering their overall excitability to adapt to persistent PTX-mediated activation, rather than from desensitization of the GABA_A_ receptor. We observed no such decrease in firing rate upon removal of DMSO (Figure S1D), indicating that the decreased excitability upon PTX withdrawal is not simply due to a change in media. Thus, we are able to measure firing rate homeostasis that occurs via adaptations in excitability in response to persistent neuronal stimulation.

Before assessing the involvement of transcription in firing rate homeostasis, we asked whether this homeostasis is cell-autonomous, that is, due to individual neurons each adapting to their individual baseline firing rates (Burrone et al., 2003; Hengen et al., 2016; Kulik et al., 2019). Alternatively, the observed firing rate homeostasis could be a non-cell-autonomous, network-level adaptation in which the firing rate of the network adapts homeostatically without firing rate homeostasis of individual neurons (Slomowitz et al., 2015). To distinguish between these possibilities, we compared the firing rates of individual neurons before and after firing rate homeostasis. We found that individual neurons have similar firing rates before and following homeostasis, as demonstrated by their clustering around the unity line in plots of post-homeostasis firing rate against the baseline firing rate (Figure 2A). In contrast, post-stimulation (i.e., 3h after PTX addition), the same neurons have increased firing rates relative to baseline (Figure 2B). Notably, as observed previously for firing rate homeostasis in response to activity blockade (Slomowitz et al., 2015), a substantial fraction of neurons do not fall exactly on the unity line, indicating that many do not return exactly to their baseline firing rates. However, this “drift” in firing rates over the course of the assay is evident in both PTX-treated neurons, which underwent firing rate homeostasis, and DMSO-treated neurons, which received no stimulation (Figures 2A,C). Indeed, the median firing rate drifts between baseline and post-homeostasis are not significantly different between DMSO- and PTX-treated neurons (Figure 2C). This equivalency between DMSO- and PTX-treated neurons indicates that the apparent failure to adapt perfectly is expected based on the observed firing rate drift, which could be due to actual fluctuation or to noise in the measurements of firing rate. In either case, our findings suggest that individual neurons homeostatically maintain their firing rates in the face of excitatory stimulation at least within a range, if not precisely.

**Figure 2.**
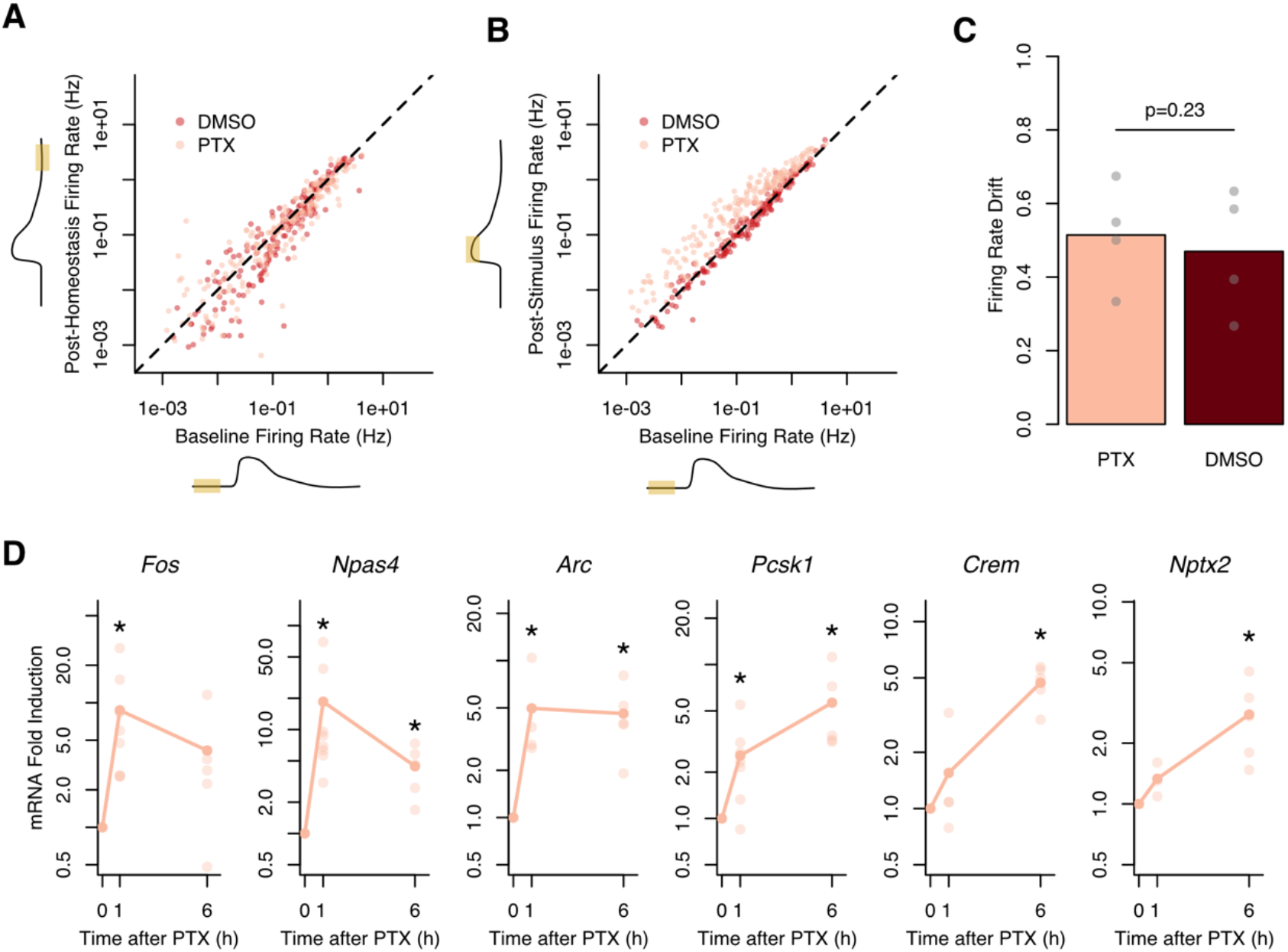
GABA_A_ blockade leads to firing rate homeostasis in individual units and induces expression of ARGs in cultured neurons. (A) A representative replicate showing a comparison of firing rates before and after homeostasis (baseline, −3h-0h; post-homeostasis, 27-30h). Each point represents an individual unit (units/replicate: DMSO, 181; PTX, 182). For PTX treatment, 42% of units showed firing rates higher in the last 3 h than at baseline; this is not greater than 50% expected: p = 0.989 binomial test. For DMSO treatment, 37% of units showed firing rates higher in the last 3 h than at baseline; this is not greater than 50% expected: p = 0.999 binomial test. (B) Same as (A) but comparing firing rates from before stimulation (baseline, −3h to 0h) to soon after addition of PTX or DMSO (post-stimulation, 0.5-3.5h). For PTX treatment, 100% of neurons showed firing rates higher in the last 3 h than at baseline; this is greater than 50% expected: p < 10^−15^ binomial test. For DMSO treatment, 46% of neurons showed firing rates higher in the last 3 h than at baseline; this is not greater than 50% expected: p = 0.883 binomial test. (C) Comparison of firing rate drift at the end of the assay (27-30h) in DMSO-treated and PTX-treated cultures. Firing rate drift was calculated as the |log2(post-homeostasis/baseline)| to weight changes from a fold change of 1 equally. n=4 biological replicates (units/replicate: DMSO, 87-187; PTX, 84-182), 1 replicate from figure 1 was eliminated because it failed to undergo firing rate homeostasis. Dots are the median fold change from individual replicates, and bar heights are the mean of the medians. P-value represents a paired t-test on median firing rate drift. (D) PTX stimulation induces mRNA expression. Fold induction calculated from Gapdh-normalized values from RT-qPCR. Each lighter dot represents the mRNA expression of a single replicate, and the darker dots show the mean expression. *p<0.006 t-test on log2(fold change), FDR<0.08. (n=3-8 biological replicates).

If transcription were required for firing rate homeostasis, we expect that PTX stimulation would induce ARG transcription. Using qPCR, we found that the same PTX stimulation used in our homeostasis assay induces transcription of activity-regulated genes in several mechanistically distinct classes (Tyssowski et al., 2018). We observed induction of the rapid primary response genes, *Arc, Fos*, and *Npas4*, the delayed primary response genes, *Pcsk1* and *Crem*, and the secondary response gene, *Nptx2* (Figure 2D). Notably, *Arc* and *Nptx2* regulate homeostatic plasticity (Chang et al., 2010; Shepherd et al., 2006). Because we observed transcription of several classes of ARGs, we expect that much of the ARG program, as identified in other experiments (Yap and Greenberg, 2018), is transcriptionally induced in neurons undergoing firing rate homeostasis in response to PTX.

### Firing rate homeostasis occurs in the absence of *Arc*

To test the requirement of transcription for firing rate homeostasis, we first focused on the requirement of the ARG Arc, as it is required for homeostatic synaptic scaling (Shepherd et al., 2006), a process that may underline firing rate homeostasis. To determine whether *Arc* is required for firing rate homeostasis, we use MEAs to measure the firing rates of cultured neurons from *Arc* knockout (KO) mice (Wang et al., 2006; Figure S3A) over 30 hours of PTX stimulation. In contrast with our prediction that *Arc* would be important for firing rate homeostasis, we found that by 27 hours of stimulation, the median firing rate and distribution of firing rates of *Arc* KO neurons return to baseline levels (Figure 3A, S3B), indicating that *Arc* KO neurons undergo firing rate homeostasis. We next asked whether *Arc* KO might delay firing rate homeostasis. We determined the time at which the median firing rate for each biological replicate returned to baseline levels after PTX addition (see Methods). This analysis revealed that the time-scale of homeostatic firing rate adaptation in *Arc* KO neurons is indistinguishable from that in their heterozygous littermates (Figure 3B), indicating that neurons lacking *Arc* undergo firing rate homeostasis with normal kinetics. Furthermore, we confirmed that compared to neurons from heterozygote littermates, *Arc* KO neurons showed similar baseline firing rate medians and distributions (Figure 3C, S3C) and responded similarly to PTX (Figure 3D). In addition, treatment with a DMSO vehicle in the place of PTX did not perturb firing rates (Figure S3D), indicating that the observed increases in firing rate are due to PTX treatment. Therefore, *Arc* KO neurons undergo firing rate homeostasis that is indistinguishable—both in extent of adaptation and kinetics—from that of control neurons, indicating that firing rate homeostasis still occurs in the absence of Arc.

**Figure 3.**
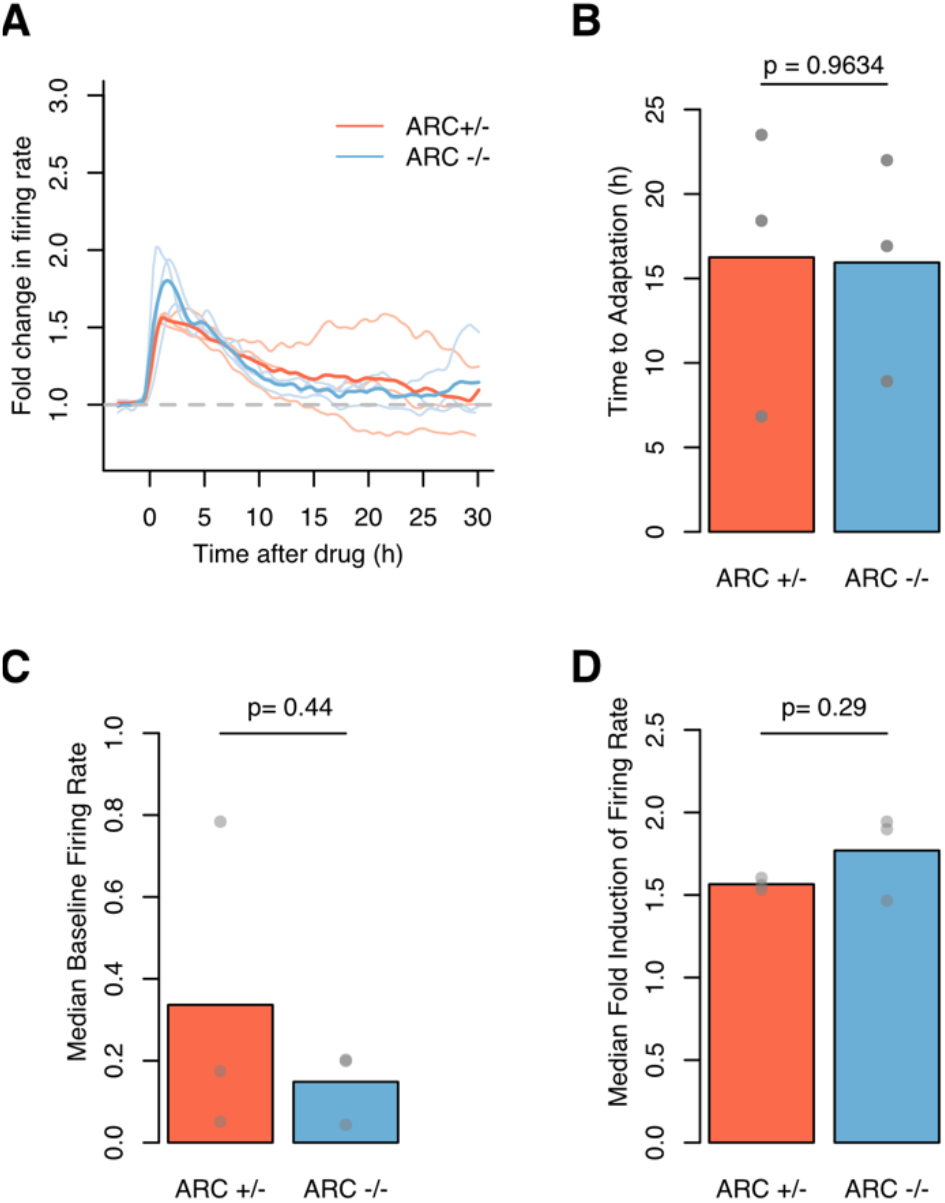
Firing rate homeostasis occurs in the absence of ARC. (A) The fold change in firing rate over the course of the assay. PTX (2.5uM) was added at time 0. Lighter lines are medians from each of n=3 biological replicates (units/replicate: Arc +/−, 64-135; Arc −/−, 20-128). Darker lines represent the mean of the median firing rates for replicates from each genotype. Arc −/− return to their baseline firing rates by the end of the assay: p = 0.46, t-test of log fold change in firing rates between baseline and last 3h of the assay (−3h to Oh, vs. 27-30h), testing difference from fold change = 1. Fold change in firing rate at the end of the assay is the same between Arc −/− and Arc +/− neurons: p =0.77, t-test on log(fold change in firing rate). (B) The time to return to baseline firing rate is indistinguishable between Arc +/− and Arc −/− neurons. Grey dots represent individual replicates (n=3). Bar height represents the mean of all replicates. P-value from a two-sided t-test. (C) Median baseline firing rates are indistinguishable between Arc +/− and Arc −/− cultures. Dots represent median firing rates before stimulation for each replicate (n=3). Bar heights represent the mean of those medians. P-value from a paired t-test on median firing rates. (D) Fold induction in firing rate upon PTX treatment is indistinguishable between Arc +/− and Arc −/− cultures. Dots represent median fold inductions of post-PTX firing rates for each replicate (n=3). Bar heights represent the mean of those medians. P-value from a paired t-sum test on log(fold change).

### Firing rate homeostasis occurs in the absence of SRF and AP1

We next hypothesized that other ARGs may compensate for the loss of Arc, or be independently required, in regulating firing rate homeostasis. We therefore aimed to simultaneously block the activity-regulated transcription of many ARGs by manipulating activity-regulated transcription factors. We focused on two transcription factors. First, we knocked out SRF, which is required for induction of a large subset of ARGs (Kuzniewska et al., 2016; Lösing et al., 2017; Ramanan et al., 2005). Second, we knocked out AP1, which regulates the slowly-induced ARGs (Vierbuchen et al., 2017; Yap and Greenberg, 2018), including *Nptx2* and *Igf1*, which regulate excitatory-inhibitory balance (Chang et al., 2010; Malik et al., 2014; Mardinly et al., 2016). We dissected cortical neurons from either an SRF conditional knockout (cKO) mouse line homozygous for floxed SRF (Ramanan et al., 2005) or an AP1 triple cKO mouse line with floxed alleles of the AP1 subunits *Fos, Fosb*, and *Junb* (Vierbuchen et al., 2017). We treated neurons from each mouse line with AAV-CaMKII-Cre to knockout SRF or AP1 in excitatory neurons and confirmed conditional KO by western blot, immunocytochemistry, or qPCR (Figures S4A, S5A). In all analyses, we compared Cre-infected neurons to neurons of the same genotype infected with AAV-CaMKII-GFP.

To determine whether SRF or AP1 are required for firing rate homeostasis, we measured the firing rates of SRF and AP1 cKO neurons in response to 30 hours of PTX stimulation. By 27h of stimulation, the median firing rate and distribution of firing rates of Cre-treated SRF and AP1 cKO neurons return to baseline levels (Figure 4A, S4B, 5A, S5B), indicating that neurons lacking SRF or AP1 still undergo firing rate homeostasis. Furthermore, the time-scale of homeostatic firing rate adaptation in Cre-treated SRF and AP1 cKO neurons is indistinguishable from that of GFP-treated neurons (Figure 4B, 5B), suggesting that neurons lacking SRF or AP1 undergo firing rate homeostasis with normal kinetics. For both genotypes, Cre- and GFP-infected neurons have indistinguishable baseline firing rates (Figure 4C, 5C, S4C, S5C) and similar responses to PTX (Figure 4D, 5D). Furthermore, treatment of Cre-infected neurons with a DMSO vehicle in the place of PTX did not perturb firing rates (Figure S4D, S5D), indicating that observed increases in firing rate are due to PTX treatment. Therefore, SRF KO and AP1 KO neurons undergo firing rate homeostasis that is indistinguishable–both in extent of adaptation and kinetics–from that of control neurons, indicating that firing rate homeostasis can stil occur in the absence of SRF or AP1.

**Figure 4.**
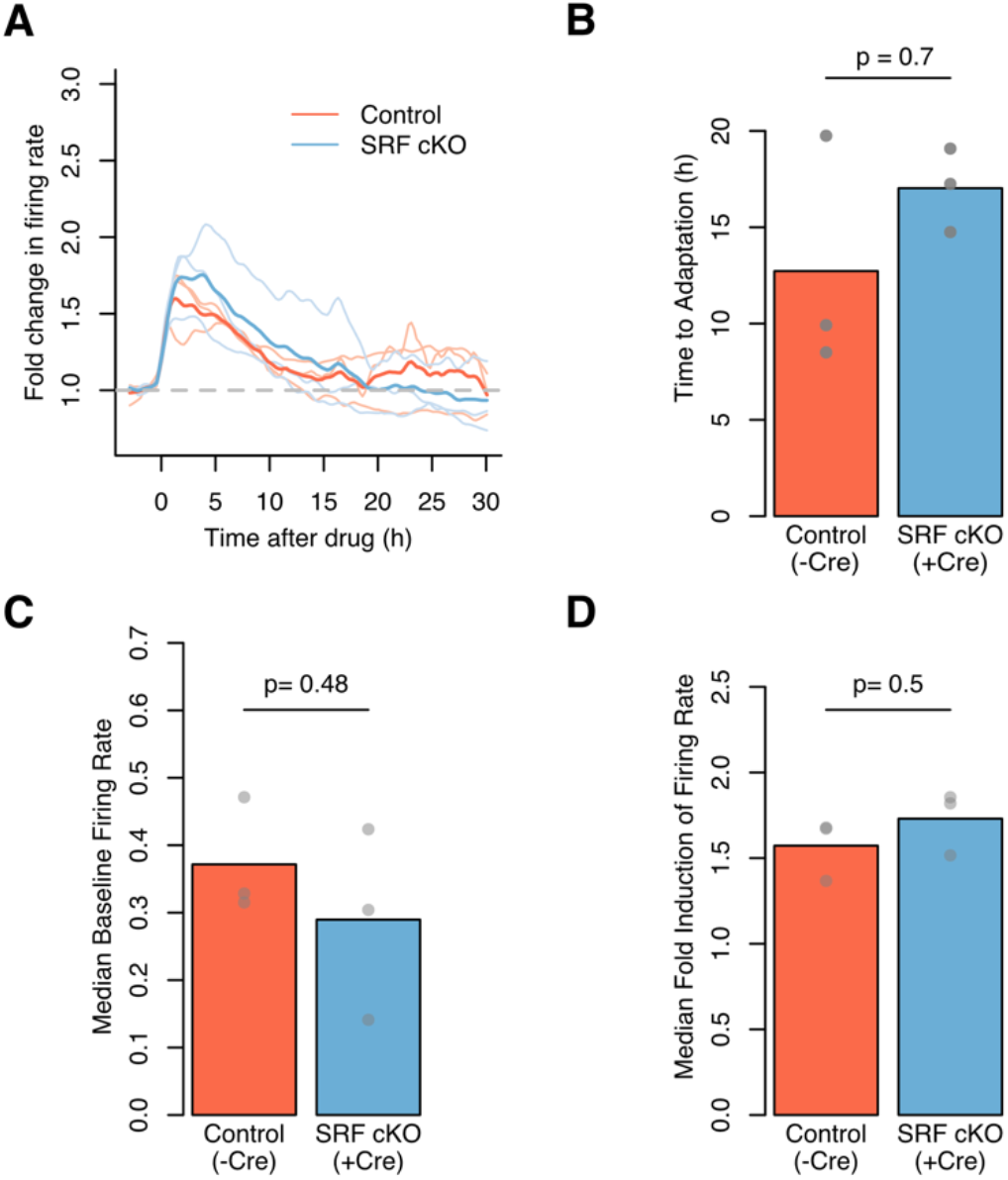
Firing rate homeostasis occurs in the absence of SRF. (A) The fold change in firing rate over the course of the assay. PTX (2.5uM) was added at time 0. Lighter lines are medians from each of n = 3 biological replicates (units/replicate: Control, 32-113; SRF KO, 26-174). Darker lines represent the mean of the median firing rates for replicates from each genotype. SRF cKO adapt back to their baseline firing rates by the end of the assay: p=0.71,t-test of log fold change in firing rates between baseline and last 3h of the assay (−3h to 0h, vs. 27-30h), testing difference from fold change = 1. Fold change in firing rate at the end of the assay is the same between SRF cKO and control neurons: p=0.43, t-test on log(fold change in firing rate). (B) The time to return to baseline firing rate is indistinguishable between control and SRF cKO neurons. Grey dots represent individual replicates (n=3). Bar height represents the mean of all replicates. P-value from a t-test. (C) SRF cKO and control median baseline firing rates are indistinguishable. Dots represent median firing rates before stimulation for each replicate (n=3). Bar heights represent the mean of those medians. P-value from a paired t-test on median firing rates. (D) Fold induction in firing rate upon PTX treatment is indistinguishable between control and SRF cKO cultures. Dots represent median post-PTX fold inductions of firing rates for each replicate (n=3). Bar heights represent the mean of those medians. P-value from a paired t-test on log(fold change).

**Figure 5.**
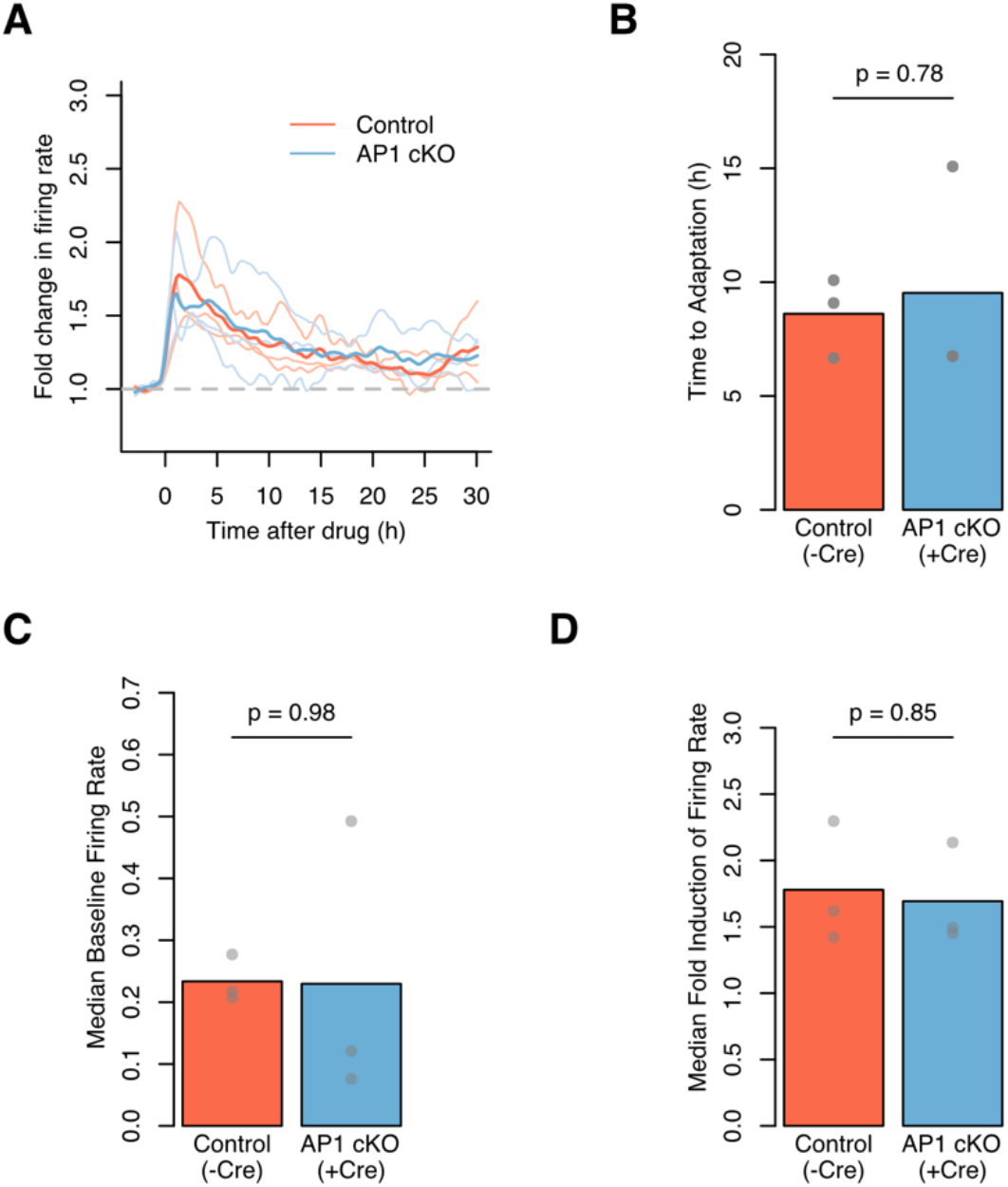
Firing rate homeostasis occurs in the absence of AP1. (A) The fold change in firing rate over the course of the assay. PTX (2.5uM) was added at time 0. Lighter lines are medians from each of n = 3 biological replicates (units/replicate: Control, 59-113; AP1 cKO: 24-67). Darker lines represent the mean of the median firing rates for replicates from each genotype. AP1 cKO adapt back to their baseline firing rates by the end of the assay: p = 0.07, t-test of log fold change in firing rates between baseline and last 3h of the assay (−3h to 0h, vs. 27-30h), testing difference from fold change = 1. Fold change in firing rate at the end of the assay is the same between AP1 cKO and control neurons: p =0.75, t-test on log(fold change in firing rate). (B) The time to return to baseline firing rate is indistinguishable between control and AP1 cKO neurons. Grey dots represent individual replicates (n=3). Bar height represents the mean of all replicates. P-value from a two-sided t-test (C) AP1 cKO and control median baseline firing rates are indistinguishable. Dots represent median firing rates before stimulation for each replicate (n=3). Bar heights represent the mean of those medians. P-value from a paired t-test on median firing rates. (D) Fold induction in firing rate upon PTX treatment is indistinguishable between control and AP1 cKO cultures. Dots represent median post-PTX fold inductions of firing rates for each replicate (n=3). Bar heights represent the mean of those medians. P-value from a paired t-sum test on log(fold change).

### Firing rate homeostasis occurs in the absence of transcription

We next sought to entirely block ARG induction to rule out the possibility that ARGs compensate for each other in regulating homeostasis. To acutely block all ARG transcription, we added the transcription inhibitor actinomycin D (ActD; 1mg/mL) to wild-type neurons 30 minutes before PTX stimulation and confirmed that ActD treatment blocked ARG induction (Figure 6A). To determine whether neurons can undergo firing rate homeostasis in the absence of transcription, we observed the firing rates of ActD-treated neurons in response to 30 hours of PTX stimulation. As we observed with ARC, SRF, and AP1 KO neurons, by 27 hours of stimulation, the median firing rates and distributions of firing rates of ActD-treated neurons return approximately to baseline levels (Figure 6B, S6A). Furthermore, the time-scale of homeostatic firing rate adaptation in Cre-treated SRF and AP1 cKO neurons is indistinguishable from that of GFP-treated neurons (Figure 6C). Neurons treated with ActD showed no difference in their response to PTX (Figure 6D), indicating similar sensitivity to PTX. However, ActD-treated neurons stimulated with a DMSO vehicle in the place of PTX showed a rapid decrease in firing rate over the first 5-10 hours following ActD treatment (Figure S6B), indicating that ActD itself affects firing rate. Indeed, we observed that the that post-homeostasis firing rates of PTX-and-ActD-treated neurons were slightly below baseline, consistent with this ActD side-effect (Figure S6C). This consistency between post-homeostasis firing rates in PTX-treated neurons and firing rates at the end of the assay in DMSO-treated neurons suggests that the ActD-treatment side effect does not impair our ability to measure firing rate homeostasis in ActD-treated neurons, rather, it changes the baseline to which neurons adapt. However, we cannot entirely rule out the possibility that ActD treatment impairs firing rate homeostasis. That said, taken with our results from ARC, AP1, and SRF KO experiments, the firing rate homeostasis we observed in ActD-treated neurons supports the conclusion that firing rate homeostasis occurs in the absence of activity-regulated transcription.

**Figure 6.**
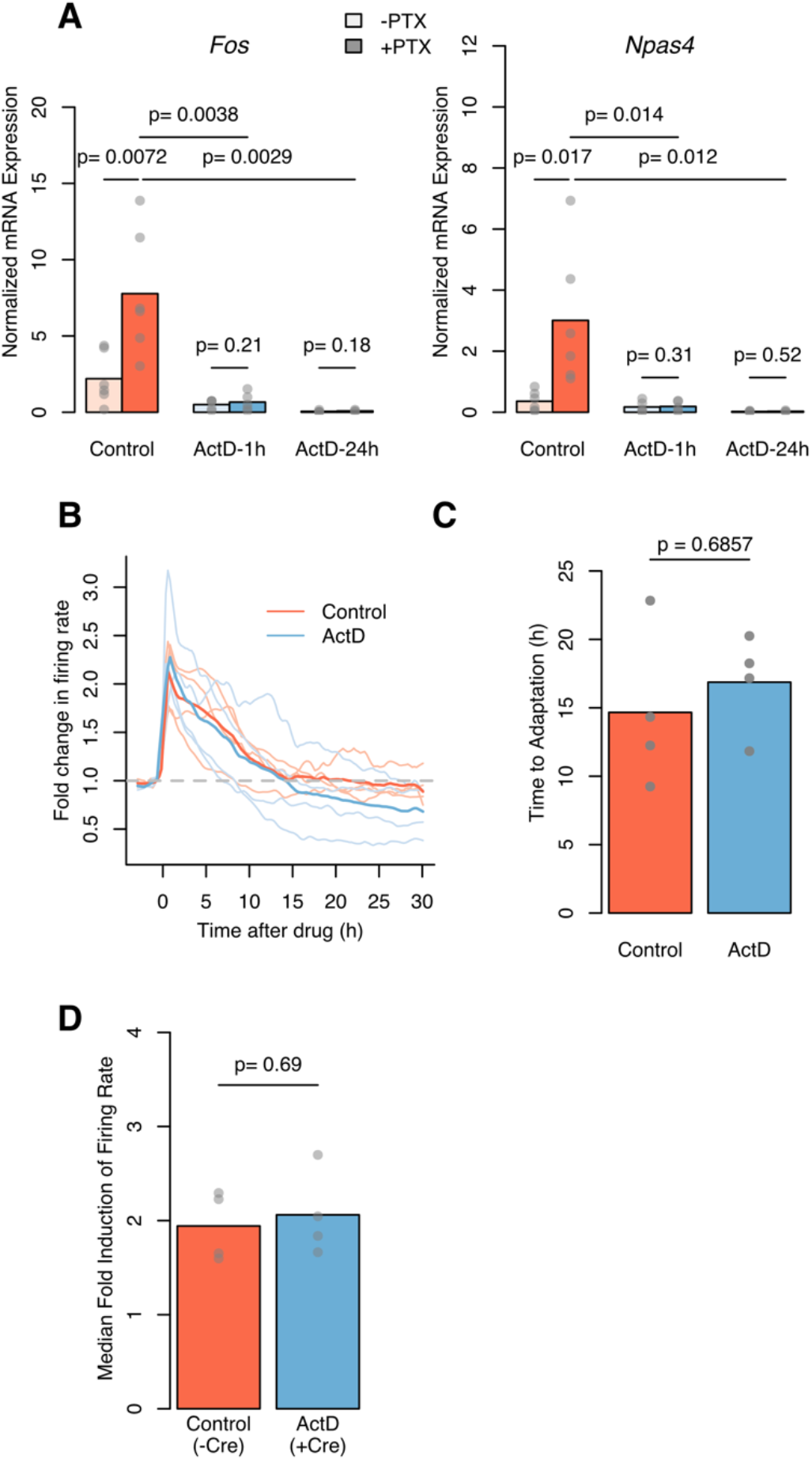
Firing rate homeostasis occurs in the presence of acute transcription blockade. (A) ActD treatment blocks activity-regulated transcription. ActD-1h samples were treated with ActD for 1.5 h and stimulated with PTX for 1h (n=6). ActD-24h samples were treated with ActD for 24.5 h and stimulated with PTX for the last lh ofActD treatment (n=3). Gapdh-normalized values from RT-qPCR. Dots represent gene expression in each biological replicate, and bars show the mean expression levels. P-values from paired, one-sided t-tests. (B) The fold change in firing rate over the course of the assay. PTX (2.5uM) was added at time 0. Lighter lines are medians from each of n=4 biological replicates (units/replicate: Control, 84-182; ActD, 58-169). Darker lines represent the mean of the median firing rates for replicates from each treatment (n=4). ActD-treated neurons adapt back to their baseline firing rates by the end of the assay: p = 0.15, t-test of log fold change in firing rates between baseline and last 3h of the assay (−3h to 0h, vs. 27-30h), testing difference from fold change = 1. Fold change in firing rate at the end of the assay is similar between ActD-treated neurons than control neurons: p =0.18, t-test on log(fold change in firing rate). (C) The time to return to baseline firing rate is indistinguishable between control and ActD-treated neurons. Grey dots represent individual replicates. Bar height represents the mean of all replicates (n=4). P-value from a two-sided t-test. (D) ActD treatment causes a slightly higher fold induction with PTX. Dots represent median post-PTX fold inductions of firing rates for each biological replicate (n=4). Bar heights represent the mean of those medians. P-value from a paired t-test on log(fold change).

## DISCUSSION

Our data indicate that neurons undergo firing rate homeostasis in the absence of activity-regulated transcription. Specifically, cultured cortical neurons from knock-out mice lacking the ARG *Arc*, or those without the activity-regulated transcription factors AP1 or SRF, homeostatically adapt their firing rates in the presence of prolonged PTX stimulation. We further found that neurons also undergo firing rate homeostasis in the presence of an acute blockade of transcription with ActD. These findings demonstrate that despite the reported roles of activity-regulated transcription in several forms of homeostatic plasticity, neuronal networks can homeostatically reduce their firing rates in response to increases in circuit activity without activity-regulated transcription.

### Mechanisms underlying firing rate homeostasis

Our finding that neurons undergo firing rate homeostasis in the absence of transcription raises the question of how firing rate homeostasis is regulated. We achieved increases in activity using GABA_A_ blockade, which induces synaptic scaling down and homeostatic decreases in intrinsic excitability (Lee and Chung, 2014; Turrigiano et al., 1998). Puzzlingly in light of our findings here, previous work suggests that both of these forms of homeostasis may be mediated by activity-regulated transcription (Cho et al., 2016; Diering et al., 2017; Goold and Nicoll, 2010; Kulik et al., 2019; O’Leary et al., 2014; Shepherd et al., 2006). Synaptic scaling down is impaired by knockout or knockdown of the ARGs Arc, *Homer1a*, and *Plk2* (Diering et al., 2017; Hu et al., 2010; Seeburg et al., 2008; Shepherd et al., 2006; Shepherd and Bear, 2011). Because we observed firing rate homeostasis in *Arc* knockout neurons and while acutely inhibiting transcription with ActD, we speculate that the firing rate homeostasis we observe may be achieved independently of synaptic scaling.

The role of transcription in regulating intrinsic excitability is less clear, and we therefore suspect that changes in intrinsic excitability could underlie the firing rate homeostasis observed here. It has been proposed that ion channel genes (e.g., potassium channels) could be transcriptionally induced in response to increases in activity and act to homeostatically decrease intrinsic excitability and thus firing rates (Cho et al., 2016; Kulik et al., 2019; O’Leary et al., 2014). However, no studies have directly linked activity-dependent transcriptional regulation of potassium channels to homeostatic regulation of intrinsic excitability. Therefore, activity-regulated-transcription-independent potassium channel regulation, such as translocation to the membrane, could underlie the firing rate homeostasis that we observe. In addition to regulation of potassium channels, neurons also decrease their intrinsic excitability by moving their axon initial segments further from the cell body (Grubb and Burrone, 2010; Kuba et al., 2010), a process that has not been suggested to require activity-regulated transcription. Thus, regulation of intrinsic excitability, either via changes in ion channel use or in axon initial segment position, remains a plausible underlying mechanism of firing rate homeostasis in response to persistent PTX treatment.

We also considered whether several other forms of homeostatic plasticity that have not yet been demonstrated to occur following GABA_A_ blockade might underlie the firing rate homeostasis we observe. One such mechanism is alteration in E/I balance, the number of excitatory relative to inhibitory synapses. Our manipulations of activity-regulated transcription likely prevent some alterations in E/I balance. Homeostatic decreases in excitatory synaptic input onto excitatory neurons (Goold and Nicoll, 2010) are blocked by acute blockade of transcription, and knock-out experiments have demonstrated that homeostatic changes in inhibitory synapse number onto specific neuronal subtypes requires several individual ARGs (Bloodgood et al., 2013; Hartzell et al., 2018; Mardinly et al., 2016; Spiegel et al., 2014). However, there are many other potentially-transcription-independent E/I balance regulation mechanisms. We thus suspect that E/I balance could regulate the firing rate homeostasis we observe.

Finally, firing rate homeostasis could be achieved via several additional forms of homeostatic plasticity, including presynaptic plasticity and alterations in Hebbian plasticity. First, the probability of presynaptic vesicle release onto excitatory neurons is altered homeostatically, and this alteration occurs in neurons that undergo firing rate homeostasis (Burrone et al., 2003). Presynaptic homeostasis can occur locally (i.e., within dendrites) rather than globally (i.e., throughout the whole neuron), making it a good candidate for ARG-independent homeostasis (Yu and Goda, 2009). In addition, neurons in our experiments may adapt their firing rates by altering their threshold for Hebbian plasticity (Bienenstock et al., 1982; Fernandes and Carvalho, 2016). However, we note that alteration in the threshold for Hebbian plasticity has been suggested, but not conclusively demonstrated, to require *Arc* (Shepherd and Bear, 2011), which is not required for firing rate homeostasis. While we do not know what cellular mechanisms underlie firing rate homeostasis in our experiments, our results demonstrate that at least one of them must be able to operate in the absence of transcriptional induction.

### Transcription-dependent homeostasis in other contexts

Although we find that neurons undergo firing rate homeostasis in the absence of activity-regulated transcription in response to persistent PTX stimulation, it is possible that a loss of activity-regulated transcription may impair firing rate homeostasis in other contexts. Indeed, different homeostatic mechanisms are engaged in response to different alterations in neuronal activity (Bridi et al., 2018; Kulik et al., 2019). Therefore, some alterations in circuit input might induce transcription-dependent mechanisms of firing rate homeostasis. For example, given that several ARGs have been implicated in regulation of inhibitory synapses onto excitatory neurons (Bloodgood et al., 2013; Gray and Spiegel, 2019; Hartzell et al., 2018; Spiegel et al., 2014), the ARG program may be more effective, and thus important, in regulating firing rate homeostasis in response to perturbations that, unlike PTX, do not themselves block inhibition onto excitatory neurons. Furthermore, neurons may have redundant mechanisms to achieve firing rate homeostasis, and thus require transcription only when other mechanisms are impaired.

Consistent with this idea, neurons can achieve the same firing rates and firing patterns with many different ion channel compositions, suggesting there are multiple ways to adjust ion channel use to achieve firing rate homeostasis (Marder and Goaillard, 2006). Therefore, activity-regulated-transcription-dependent mechanisms of achieving firing rate homeostasis could be operating in our assay, but neurons may be able to compensate for their loss with transcription-independent mechanisms.

It is also possible that we failed to observe a role for transcription in regulating firing rate homeostasis because firing rate is not the activity parameter that transcription-dependent mechanisms of homeostatic plasticity regulate. For example, criticality is a network-level property of neuronal circuits that describes circuits that have stable activity, that which neither dies out nor increases over time (Shew and Plenz, 2013). A recent study demonstrated that in response to visual deprivation, the mouse visual cortex initially enters a subcritical state, but returns to criticality over a period of 48h, faster than the cortex undergoes firing rate homeostasis in response to the same perturbation (Ma et al., 2018). Modeling suggests that homeostasis of criticality could be mediated by transcription-dependent synaptic scaling, raising the possibility that transcription-dependent homeostatic plasticity controls criticality rather than firing rate. In addition to firing rate and criticality, neurons also undergo homeostasis of firing pattern (Marder and Goaillard, 2006), and transcription could also potentially regulate firing pattern homeostasis, for example through regulation of ion channel expression. Our work thus does not rule out a role for activity-regulated transcription in neuronal activity homeostasis, but raises the possibility that to identify its role, it may be important to consider parameters of neuronal activity other than firing rate.

## ACKNOWLEDGEMENTS

We thank Yasmin Escobedo Lozoya and members of the Gray lab for feedback throughout the project. KMT received funding from the NSF Graduate Research Fellowship Program DGE1144152, DEG1745303. JMG’s lab is supported by NIHMH116223, the Harvard-Armenise Foundation, and the Kaneb family.

## EXTENDED DATA FIGURES/TABLES

**Figure S1.**
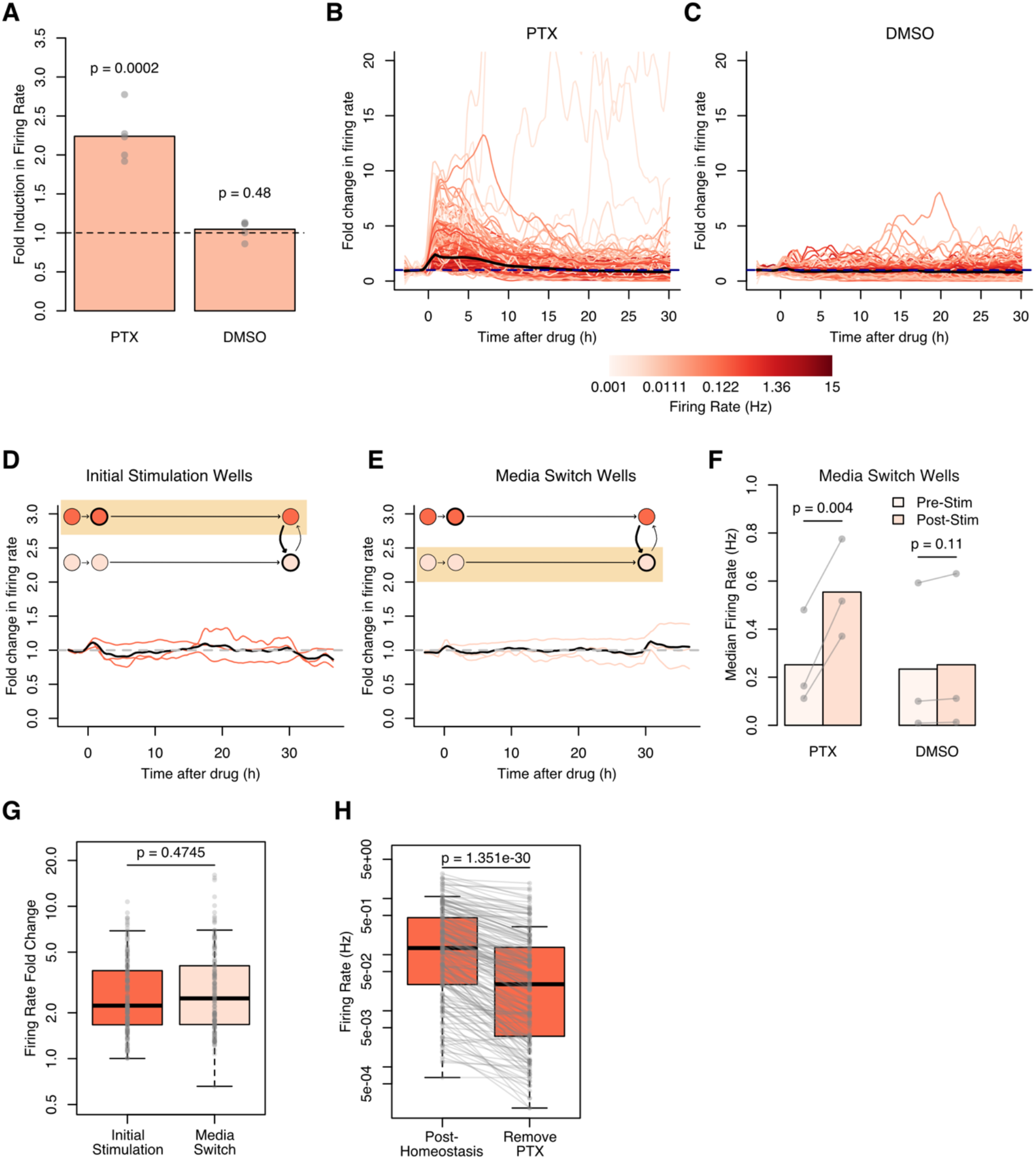
Cultured neurons homeostatically adapt their firing rates in response to GABA_A_ blockade. (A) Fold induction in firing rates after PTX or DMSO addition (0.3-3.5h/0-3h), n=5 biological replicates each. Dots represent the median fold induction from all neurons in each replicate. Bar heights represent the mean of the medians. P-values from t-tests on median log fold inductions, testing a difference from a fold induction of 1. (units/replicate: DMSO, 77-187, PTX, 31-182). (B) Representative example of the fold change in firing rate for one replicate (n=182 units). PTX (2.5uM) was added at time 0. Colored lines are the fold changes in firing rates of individual neurons, colored by their mean baseline firing rate in Hz. The black line represents the median firing rate for the replicate. (C) Same as (B) but DMSO was added at time 0 instead of PTX (n=181 units). (D) The fold change in firing rate over the course of the media switch assay in the wells initially treated with DMSO. DMSO was added at 0h and media changed (i.e., DMSO removed) at 30h. Colored lines are medians from individual replicates (n=3 biological replicates; 81-187 units/replicate). The black line represents the mean median firing rate. (E) The fold change in firing rate over the course of the media switch assay in the wells that received DMSO upon media switch. DMSO from initially-treated wells was added at 30h. Colored lines are medians from individual replicates (n=3 biological replicates, 67-214 units/replicate). Black line represents the mean median firing rate. (F) Quantification of firing rate responses to media switch in panels 1I and S1E. Dots represent the median firing rate over a 3h-period for each replicate either before the media switch (pre-Stim) or after (post-Stim). Bar heights represent the mean of the medians. P-values from a one-sided t-test on the medians of firing rates. (G) A representative replicate showing the fold change in firing rates upon PTX addition to initially-stimulated wells (n=182 units) and media switch wells (n=159 units). Firing rates used to calculate fold change are the mean firing rate over a 3h period. P-value from a rank-sum test. (H) A representative replicate showing the decrease in firing rate upon PTX removal (n=182 units). Firing rates are the mean firing rate of an individual neuron over a 3h period. P-value from a paired rank-sum test. p<10^−13^ for all replicates.

**Figure S3.**
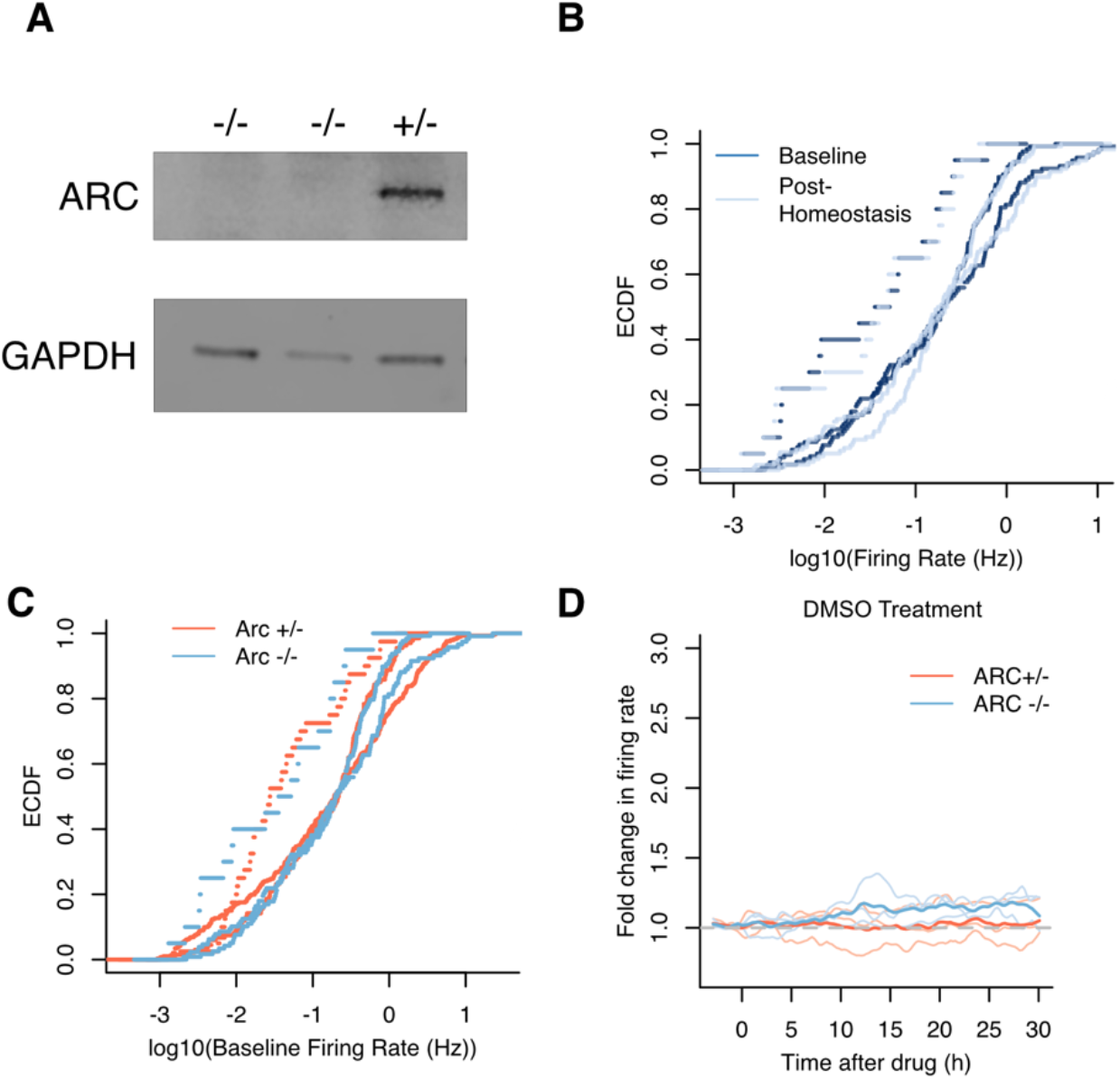
Firing rate homeostasis occurs in the absence of ARC. (A) Confirmation of ARC KO. Cultured neurons were stimulated with potassium chloride for 2h to induce ARC protein before collecting. A representative image from one of n=3 biological replicates. (B) Empirical cumulative distributions of firing rates of Arc −/− neurons before adding PTX (−3-0h, baseline) and at the end of the recording (27-30h, post-homeostasis). The value for each neuron is the mean firing rate over a 3h period. For all replicates, the distribution of firing rates is not significantly different between baseline and post-homeostasis (Kolmogorov-Smirnov test, p = 0.98, 0.17, 0.99 for each replicate, n=3 biological replicates, 20-128 unitsIreplicate). (C) Empirical cumulative distributions of firing rates before adding PTX (baseline) of Arc +/− and Arc −/− neurons. The value for each neuron is the mean firing rate over a 3h period. The distributions of Arc −/− and Arc +/− neurons were compared for n=3 biological replicates (unitsIreplicate: Arc −/−, 20-128; Arc +/− 64-135), p=0.51,0.24,0.99 for each replicate (Kolmogorov-Smirnov test). (D) The fold change in firing rate over the course of the assay when DMSO was added at time 0 instead of PTX. Lighter lines are medians from individual replicates (n=3 replicates; unitsIreplicate: Arc +/−123-215; Arc −/−, 40-291). Darker lines represent the mean of the median firing rate for replicates from each genotype.

**Figure S4.**
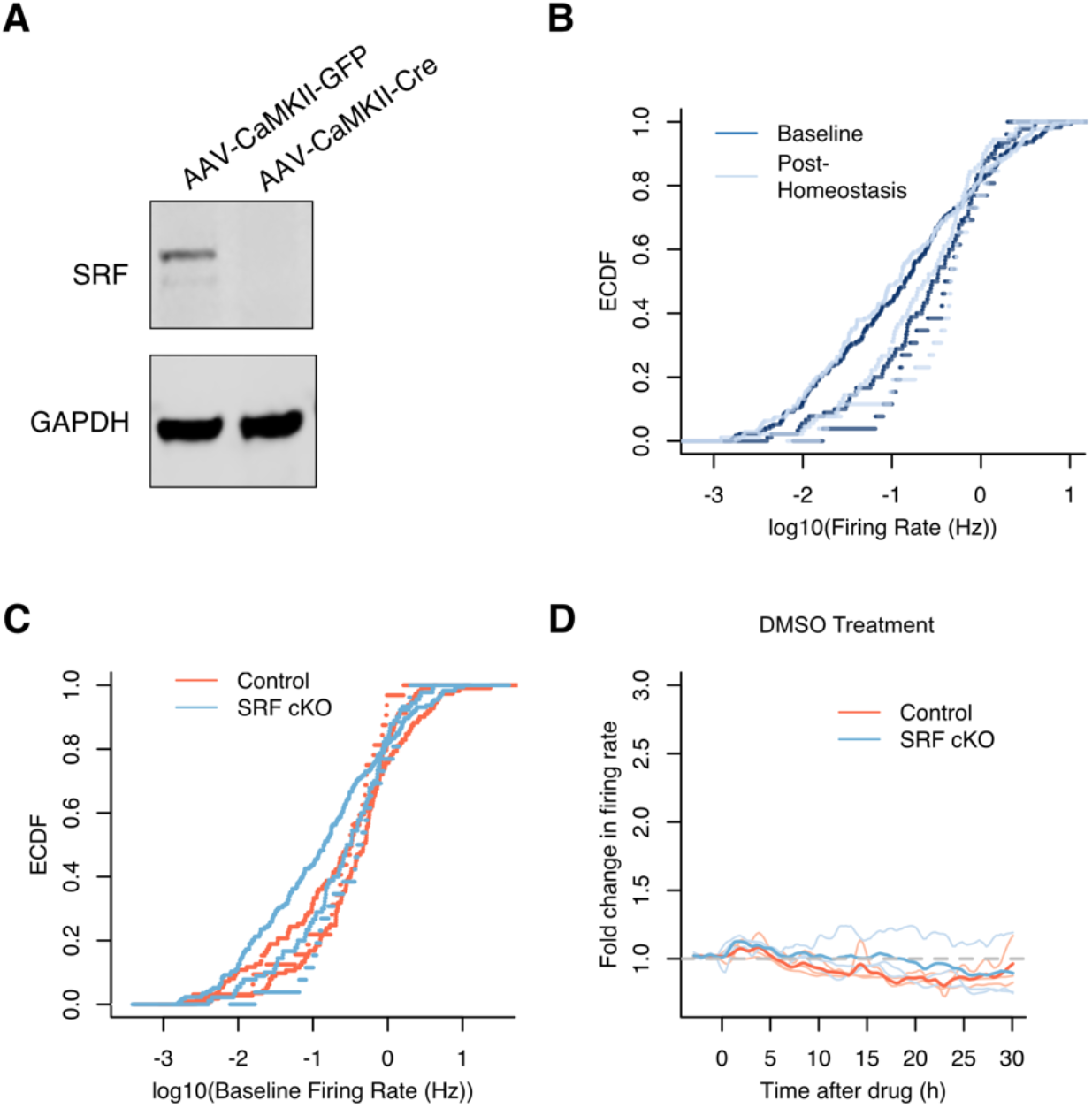
Firing rate homeostasis occurs in the absence of SRF. (A) Confirmation of SRF KO. Neurons were treated with Cre or GFP virus 14-16 days before collection. A representative image from one of n=3 biological replicates. (B) Empirical cumulative distributions of SRF cKO firing rates before adding PTX (−3-0h, baseline) and at the end of the recording (27-30h, post-homeostasis). The value for each neuron is the mean firing rate over a 3h period. For all replicates, the distribution of firing rates is not significantly different between the baseline and post-homeostasis (Kolmogorov-Smirnov test, p = 0.72, 0.93, 0.51 for each replicate, n=3 biological replicates, 130-291 units/replicate). (C) Empirical cumulative distributions of firing rates before adding PTX (baseline) of control and SRF cKO neurons. The value for each neuron is the mean firing rate over a 3h period. The distributions of control and SRF neurons were compared for n=3 biological replicates (units/replicate: Control, 32-113; SRF cKO, 26-174), p = 0.03, 0.54, 0.20 for each replicate (Kolmogorov-Smirnov test). (D) The fold change in firing rate over the course of the assay when DMSO was added at time 0 instead of PTX. Lighter lines are medians from each of n=3 biological replicates (units/replicate: Control, 151-295; SRF cKO, 130-291). Darker lines represent the mean of the median firing rates for replicates from each genotype.

**Figure S5.**
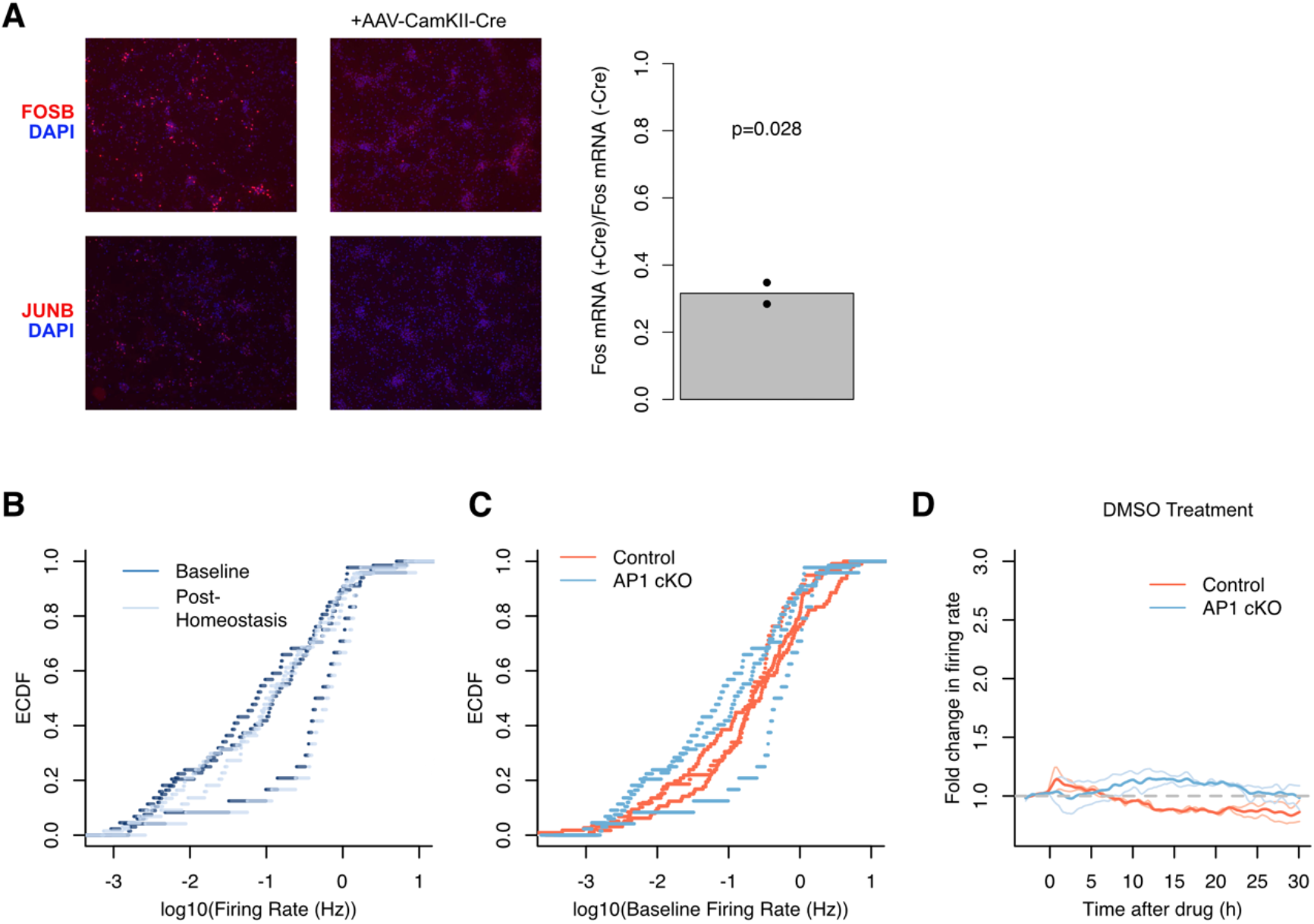
Firing rate homeostasis occurs in the absence of AP1. (A) Confirmation that Cre treatment successfully knocks out Fos, Fosb, and Junb. Immunocytochemistry for FOSB and JUNB is a representative image of n=2 biological replicates. The Fos mRNA expression was measured by qPCR on RNA from neuronal cultures exposed to 1h of PTX treatment. Bar height is the average expression and dots are individual replicates (n=2). P-value from a one-sided t-test. (B) Empirical cumulative distributions of AP1 cKO firing rates before adding PTX (−3-0h, baseline) and at the end of the recording (27-30h, post-homeostasis). The value for each neuron is the mean firing rate over a 3h period. For all replicates, the distribution of firing rates is not significantly different between the baseline and post-homeostasis (Kolmogorov-Smirnov test, p=0.64, 0.91, 0.99 for each replicate, n = 3 biological replicates, 24-67 units/replicate). (C) Empirical cumulative distributions of firing rates before adding PTX (baseline) of Control and AP1 cKO neurons. The value for each neuron is the mean firing rate over a 3h period. The distributions of control and AP1 ckO neurons were compared for n=3 biological replicates (units/replicate: Control, 59-113; AP1 cKO, 24-67), p = 0.061, 0.185, 0.193 (Kolmogorov-Smirnov test). (D) The fold change in firing rate over the course of the assay when DMSO was added at time 0 instead of PTX. Lighter lines are medians from individual replicates (n = 2 biological replicates; units/replicate: AP1 cKO, 107-117; Control, 84-119). Darker lines represent the mean of the median firing rates for replicates from each genotype.

**Figure S6.**
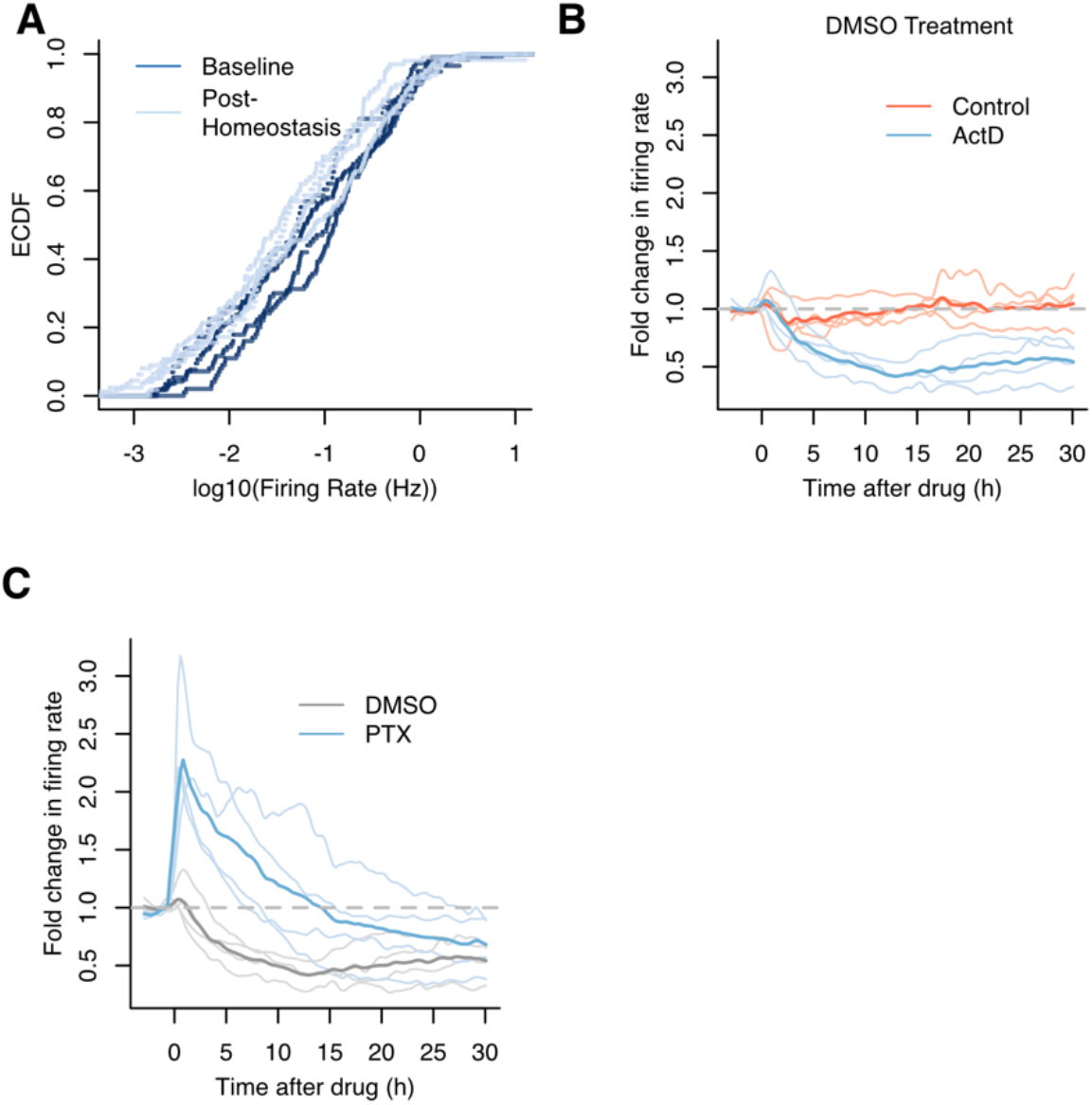
Firing rate homeostasis occurs in the presence of acute transcription blockade. (A) Empirical cumulative distributions of ActD-treated neurons’ firing rates before adding PTX (−3-0h, baseline) and at the end of the recording (27-30h, post-homeostasis). The value for each neuron is the mean firing rate over a 3h period. For each replicate, we tested the difference between the distribution of firing at baseline and post-PTX (Kolmogorov-Smirnov test, p= 0.79, 0.24, 0.004, 0.002). (n=4 biological replicates, 58-169 units/replicate). (B) The fold change in firing rate over the course of the assay when DMSO was added at time 0 instead of PTX. Lighter lines are medians from individual replicates (n=4 biological replicates; Control, 87-187; ActD, 148-214). Darker lines represent the mean of the median firing rates for replicates from each genotype. (C) The fold change in firing rate over the course of the assay when ActD was added at −0.5h, and then either DMSO or PTX was added at time 0. Lighter lines are medians from individual replicates (n=4 biological replicates; DMSO, 148-214; PTX, 58-169). Darker lines represent the mean of the median firing rates for replicates from each genotype.

**Table S1.** Alternative Stimulations Attempted.

| Drug | Mechanism of Action | Concentration | Results |
| --- | --- | --- | --- |
| Dendrotoxin | Blocks K<sup>+</sup> channels (K<sub>v</sub>1.1, K<sub>v</sub>1.2, K<sub>v</sub>1.6) | 10nM, 100nM | Increases FR, but adaptation is inconsistent |
| 4-Aminopyridine | Blocks K<sup>+</sup> channels (K<sub>v</sub>1.1, K<sub>v</sub>1.2) | 10-100 μM | Increases FR, but adaptation is inconsistent |
| Barium | Activates calcium-dependent K<sup>+</sup> current, and blocks voltage-dependent | 0.1mM | Increases FR, but does not adapt |
|  | K<sup>+</sup> current and K<sup>+</sup> leak currents |  |  |
| KCl | Increases extracellular K<sup>+</sup> | 0.1mM | Does not consistently increase FR |
| Sodium | Increases extracellular Na<sup>+</sup> | 60mM | Does not increase FR |
| OD1 | Activates Na<sup>+</sup> channels (Na<sub>v</sub>1.6, Na<sub>v</sub>1.7) | 10-300 nM | Does not increase FR |
| Bicuculline | GABA(A) antagonist | 2-40uM | Increases FR, but duration and magnitude of the increase is variable |
| Cyclothiazide | AMPA receptor agonist | 50uM, 100uM | Drug loses potency in 0.5 days |

**Table S2.**
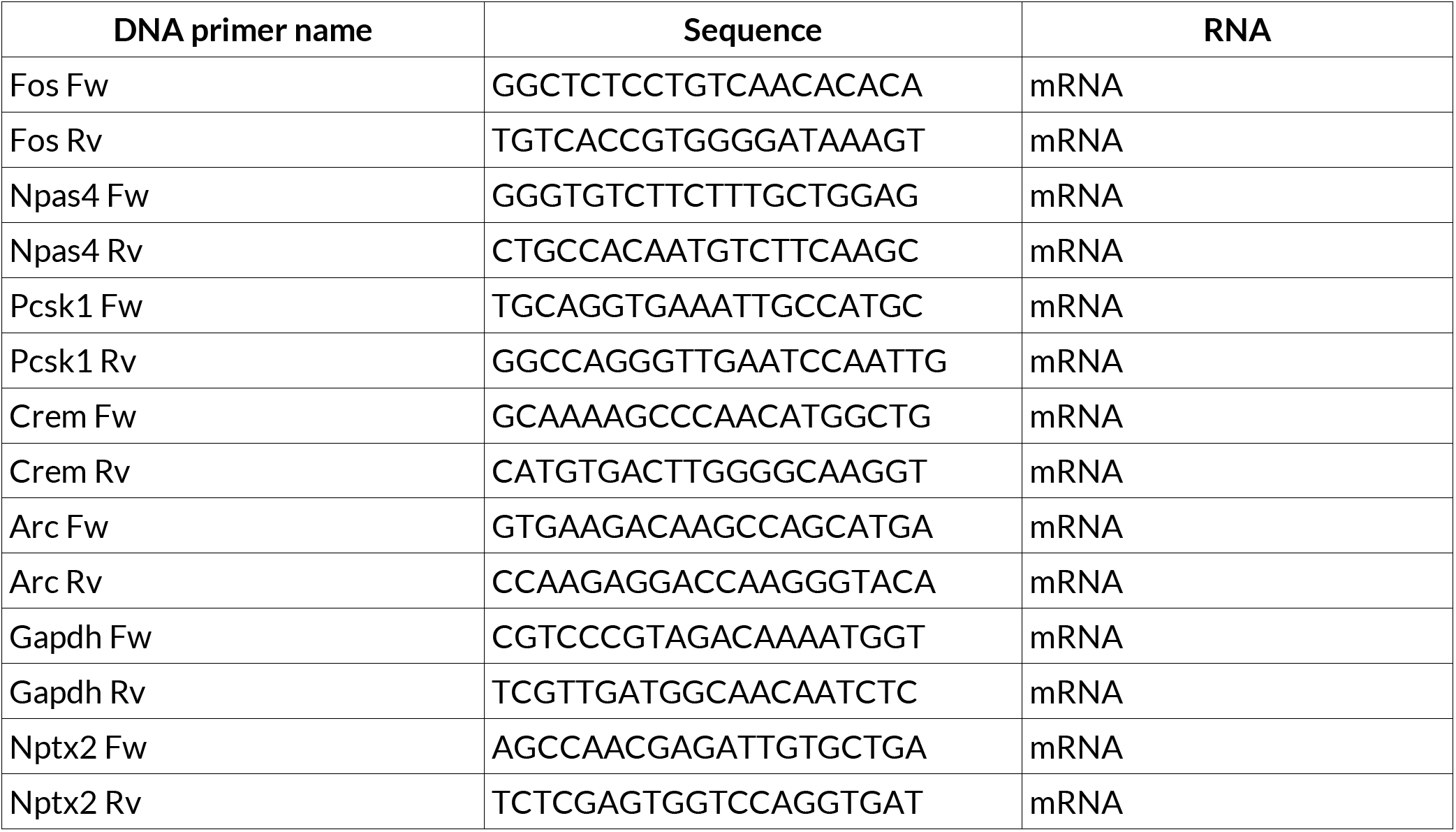
qPCR primers.

**Table S3. Data related to Figures 1 A-F, Figure 2**

Table of binned spikes for each unit of each replicate and treatment condition used to make Figures 1 A-F and Figure 2. Rows are each a 5-min time bin and columns are a unit.

**Table S4. Data related to Figures 1 G-L**

Table of binned spikes for each unit of each replicate and treatment condition used to make Figures 1 G-L. Rows are each a 5-min time bin and columns are a unit.

**Table S5. Data related to Figure 3 (Arc KO)**

Table of binned spikes for each unit of each replicate and treatment condition used to make Figure 3. Rows are each a 5-min time bin and columns are a unit.

**Table S6. Data related to Figure 4 (SRF cKO)**

Table of binned spikes for each unit of each replicate and treatment condition used to make Figures 4 Rows are each a 5-min time bin and columns are a unit.

**Table S7. Data related to Figure 5 (AP1 cKO)**

Table of binned spikes for each unit of each replicate and treatment condition used to make Figure 5. Rows are each a 5-min time bin and columns are a unit.

**Table S8. Data related to Figure 6 (ActD)**

Table of binned spikes for each unit of each replicate and treatment condition used to make Figure 6. Rows are each a 5-min time bin and columns are a unit. Data for controls (no ActD) same as data in Table S3.

